# Experimentally engineered mutations in a ubiquitin hydrolase, UBP-1, modulate *in vivo* resistance to artemisinin and chloroquine in *Plasmodium berghei*

**DOI:** 10.1101/2019.12.12.874990

**Authors:** Nelson V. Simwela, Katie R. Hughes, A. Brett Roberts, Michael T. Rennie, Michael P. Barrett, Andrew P. Waters

**Author notes:** Corresponding author: Andrew P. Waters.

## Abstract

As resistance to artemisinins (current frontline drugs in malaria treatment) emerges in south East Asia (SEA), there is an urgent need to identify the genetic determinants and understand the molecular mechanisms underpinning such resistance. Such insights could lead to prospective interventions to contain resistance and prevent the eventual spread to other malaria endemic regions. Artemisinin reduced susceptibility in SEA has been primarily linked to mutations in *P. falciparum* Kelch13, which is currently widely recognised as a molecular marker of artemisinin resistance. However, 2 mutations in a ubiquitin hydrolase, UBP-1, have been previously associated with artemisinin resistance in a rodent model of malaria and some cases of UBP-1 mutation variants associating with artemisinin treatment failure have been reported in Africa and SEA. Here, CRISPR-Cas9 genome editing and pre-emptive drug pressures was used to test these artemisinin resistance associated mutations in UBP-1 in *P. berghei* sensitive lines *in vivo*. The data demonstrate that the V2721F UBP-1 mutation results in artemisinin resistance and some low-level resistance to chloroquine, while the V2752F mutation results in high-level resistance to chloroquine and moderate resistance to artemisinins. Genetic reversal of the V2752F mutation restored chloroquine sensitivity in these mutant lines while simultaneous introduction of both mutations could not be achieved and appears to be lethal. Interestingly, these mutations carry a detrimental growth defect, which would possibly explain their lack of expansion in natural infection settings. This is the first independent, direct experimental evidence on the role of UBP-1 in artemisinin and chloroquine resistance under *in vivo* conditions.

## Introduction

Artemisinins (ARTs) in artemisinin combinational therapies (ACTs) remain the mainstay of malaria treatment globally and thus far remain mostly effective in sub-Saharan Africa where most of the disease burden occurs (WHO, 2018). However, ART (and even ACT) resistance has emerged in SEA with a risk of spreading which is seriously threatening recent gains achieved in malaria control (Hamilton et al., 2019, Dondorp et al., 2009). ART resistance is thought to be primarily conferred by specific mutations in the *Plasmodium falciparum* Kelch 13 (PfKelch13) gene, and such mutations are currently almost endemic in most parts of SEA (Mbengue et al., 2015, Ashley et al., 2014, WHO, 2018). Phenotypically, these mutations are associated with delayed parasite clearance rates *in vivo* and reduced susceptibility of ring stage parasites *in vitro* in ring stage survival assays (RSA) (Witkowski et al., 2013, Dondorp et al., 2009). Interestingly, the prevalence of PfKelch13 mutations remains low outside SEA (Menard et al., 2016) where the few observed PfKelch13 polymorphisms in sub-Saharan Africa do not associate with treatment failure and/or delayed parasite clearance rates (Sutherland et al., 2017). Moreover, large-scale genome wide association studies have revealed that polymorphisms in other genes such as multidrug resistance protein 2, ferredoxin and others also associate with delayed parasite clearance rates in SEA (Miotto et al., 2015). More recently, mutations in an independent gene, *P. falciparum* coronin (PfCoronin) have been shown to confer enhanced survival of ring stage parasites to dihydroartemisinin (DHA) (Demas et al., 2018). Deconvoluting the geographic complexities of ART resistance, genetic determinants and molecular mechanism involved would thus provide an avenue to contain or rescue the emergent ART resistance through efficient surveillance and/or suitable combinational therapies.

Mutations in a ubiquitin hydrolase, UBP-1 (HAUSP or USP7 close homologue), were previously identified to confer ART and chloroquine (CQ) resistance in the rodent infectious malaria parasite, *Plasmodium chabaudi*, after sequential experimental evolution and selection with a series of antimalarial drugs (Hunt et al., 2007). The drug resistance phenotypes that emerged resulted from *in vivo* passage and exposure of *P. chabaudi* drug sensitive AS line to sub-lethal doses of pyrimethamine, CQ, mefloquine and ARTs (Hunt et al., 2010, Hunt et al., 2007, Afonso et al., 2006). Interestingly, in these *P. chabaudi* lineages, CQ resistance at 15mg/kg emerged first and from this uncloned line, whole genome sequencing revealed 2 UBP-1 mutations (V2697F and V2728F) which were associated with the resistance phenomenon (Henriques et al., 2013, Afonso et al., 2006). Further selection of this uncloned CQ resistant line generated lines with different drug resistant profiles: 1) a line resistant to 15mg/kg mefloquine 2) a line resistant to CQ at 30mg/kg 3) a line resistant to up to 300mg/kg ART which was selected from the CQ 30mg/kg resistant line 4) a line resistant to up to 60mg/kg artesunate. Upon further cloning and genome sequencing of these lines, it was found that the UBP-1 V2728F mutation was common in the ART, CQ (30mg/kg) and mefloquine resistant lines while the V2697F mutation only fixated upon artesunate selection (Henriques et al., 2013, Hunt et al., 2010, Hunt et al., 2007). Due to the complexity of the selection procedure with multiple drugs, it has been difficult to confidently associate these UBP-1 mutations with ART and CQ resistance in the absence of appropriate reverse genetics approaches. More interestingly, UBP-1 mutation variants have been associated with ARTs decreased effectiveness in Africa and some parts of Asia (Henriques et al., 2014, Adams et al., 2018, Cerqueira et al., 2017, Borrmann et al., 2013).

In our present study, we have successfully engineered UBP-1 candidate mutations in an independent rodent model, *P. berghei* using a CRISPR-Cas9 genome editing system. We provide a direct causal link to the ART and CQ resistance profiles of these mutant lines both *in vitro* and *in vivo*. We have also characterised their relative fitness as compared to the wildtype non-mutant parasites.

## Materials and methods

### CRISPR Cas9 generation of UBP-1 mutant lines

#### Primary vectors

The Cas9 expressing plasmid ABR099 was used for targeted nucleotide replacement at the UBP-1 locus. ABR099 (Figure 1A) contains the Cas9 endonuclease driven by the Pb Ef-1α promoter, a Cas9 binding scaffold, a site for cloning the guide RNA (sgRNA) driven by the *Plasmodium yoelii* U6 promoter, an hdhfr cassette (for pyrimethamine drug resistance selection) and a linker site for insertion of homologous repair templates. sgRNAs targeting the UBP-1 locus were designed using the web based eukaryotic pathogen CRISPR guide RNA/DNA design tool (http://grna.ctegd.uga.edu/) (Peng and Tarleton, 2015) by directly inputting the sequence of interest. Primary vectors containing the sgRNA of interest were generated by annealing oligonucleotide pairs (GU4788+GU4789 and GU5206+GU5207, supplementary Table 1) encoding the guide sequence and cloning them in the dual *Esp3I* sites upstream of the Cas9 binding domain of the vector ABR099. These plasmids were called pG944 and pG960 for the GU4788+GU4789 and GU5206+GU5207 annealed guides respectively.

**Figure 1:**
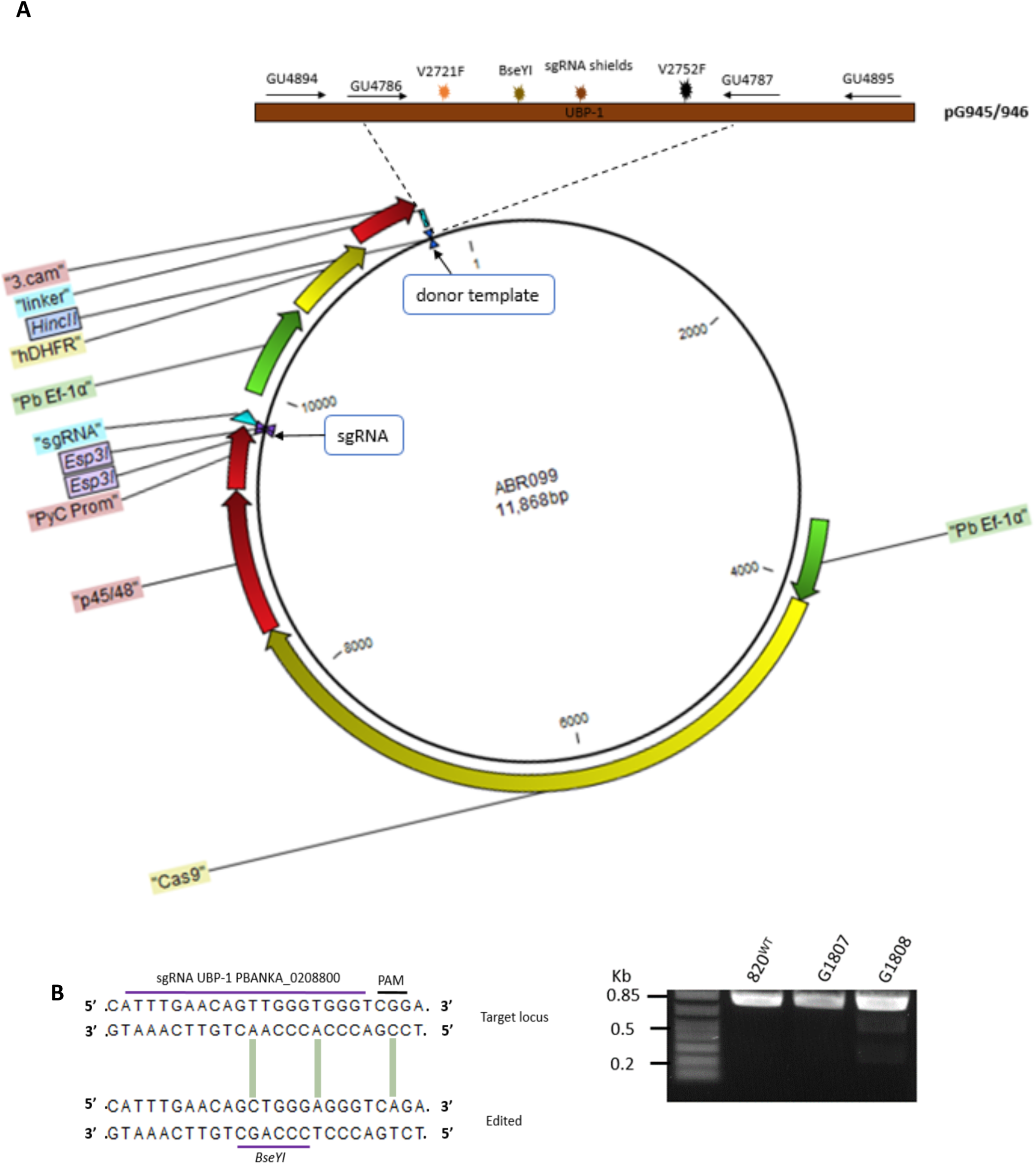
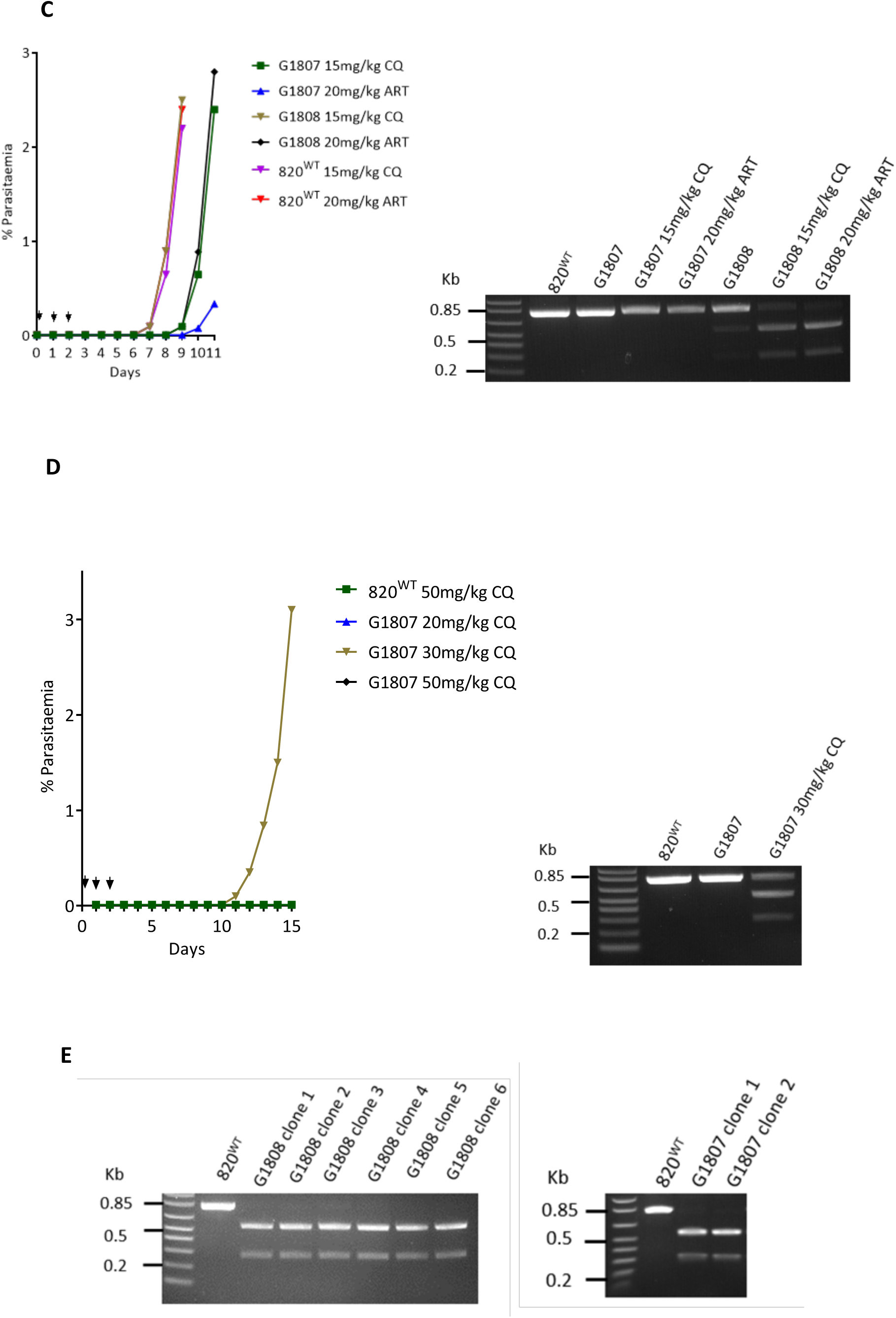
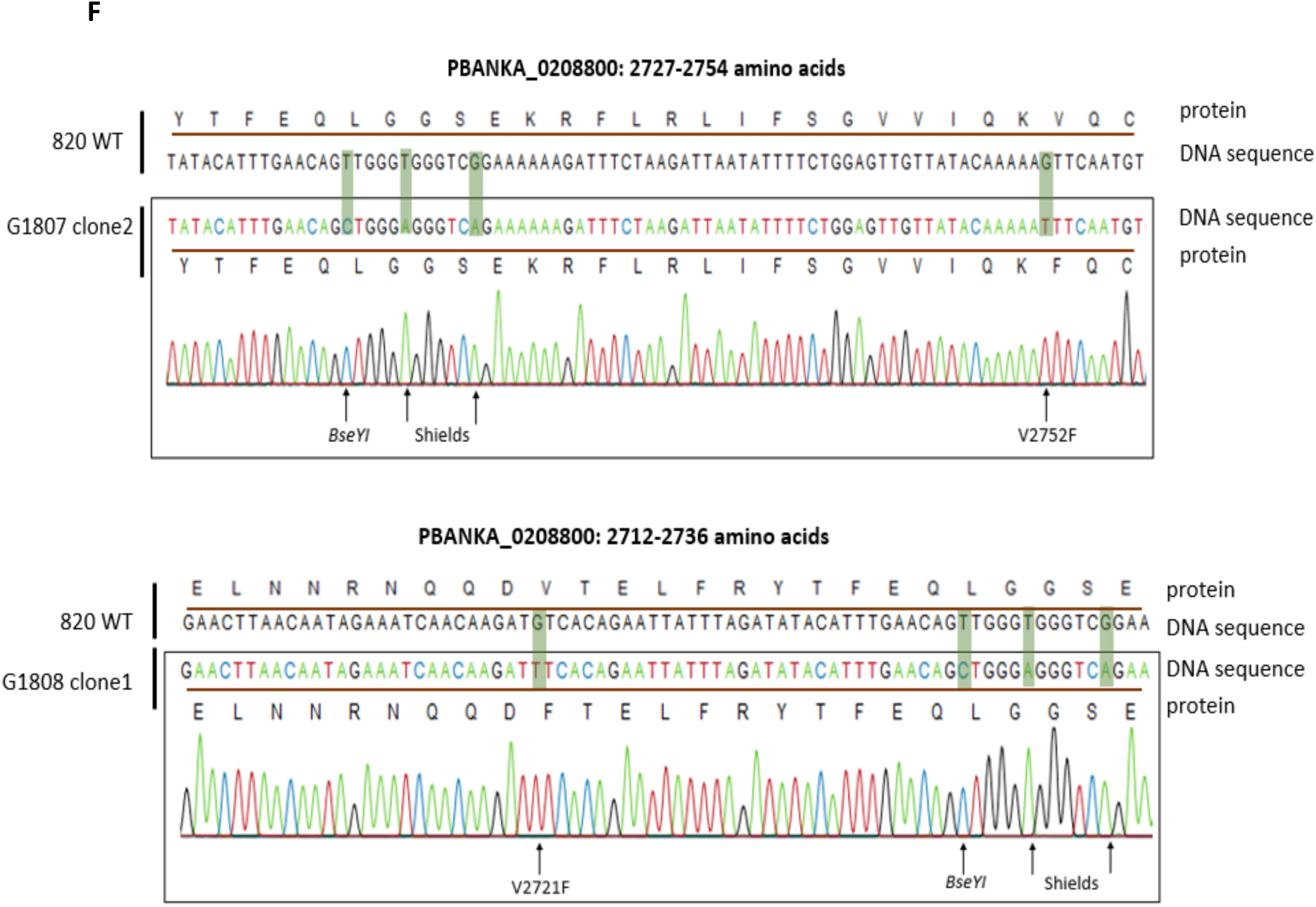
Introduction of UBP-1 mutations in *P. berghei*. **A.** Schematic plasmid constructs for the UBP-1 targeted gene editing to introduce the V2721F and V2752F mutation. The plasmid contains Cas9 and hdhfr (for pyrimethamine drug selection) under the control of the Pb EF-1α promoter and the sgRNA expression cassettes under the control of PyU6 promoter. A 20bp guide RNA was designed and cloned into sgRNA section of the vector illustrated in **A**. The donor UBP-1 sequence (516bp) is identical to the wildtype sequence albeit with the desired mutations of interest as indicated by coloured star symbols: V2752F (pG945), V2721F V2752F (pG946) and silent mutations that mutate the Cas9 binding site as well as introduce the restriction site *BseYI* for RFLP analysis. **B.** Illustrated 20bp sgRNA and RFLP analysis of mutant parasites. Successful editing in the transfected parasites was observed on day 12 after transfection and pyrimethamine drug selection. RFLP (*BseYI* digestion) analysis of the transformed lines PCR products (primers GU4894 + GU4895, 713bp) revealed <1% and ~20% efficiency for the G1807 and G1808 lines respectively as indicated by 2 distinct bands (442bp, 271bp) as compared to 713bp bands in the parent 820 line. **C.** pre-emptive challenge of the G1807 and G1808 lines with ART and CQ at 20mg/kg and 15mg/kg respectively and RFLP analysis of recrudescent parasites. Mice were infected with ~2 × 10^7 parasites IP on day 0. Treatment was started ~4 hours post infection by IP for three consecutive days. Parasitaemia was monitored by microscopy analysis until recrudescence was observed. **D.** Pre-emptive challenge of the G1807 line with higher doses of CQ and RFLP (*BseYI* digestion) analysis of the G1807 recrudescent population after challenge with 30mg/kg CQ. **E.** RFLP analysis of the cloned G1808 and G1807 ART and CQ challenged recrudescent parasites. **F.** DNA sequencing confirming successful nucleotide editing for the G1807 clone2 and G1808 clone1 line. The top sequence represents the 820^WT^ unedited sequence with positions for sgRNA, protospacer adjacent motif (PAM) and V2721F or V2752F mutations indicated. The bottom sequence illustrates the nucleotide replacements at V2721F or V2752F mutation locus and silent mutations to prevent Cas9 retargeting as well as introduce the *BseYI* restriction site for *RFLP* analysis in the G1807^V2752F^ and G1808^V2721F^ lines.

#### Mutagenesis and generation of secondary vectors

To generate the final vectors for editing the UBP-1 locus, 516bp of UBP-1 donor DNA (PBANKA_0208800) was PCR amplified using primers GU4786 and GU4787 (supplementary Table 1) designed to contain a *HincII* site at the 5’ end. The PCR product was purified, A-tailed and cloned into the TOPO 2.1 vector using the TOPO TA cloning kit (Invitrogen) according to manufacturer’s instructions. To mutate the UBP-1 locus, 3 primer sets (supplementary Table 1) complementary to the amplified UBP-1 PCR product were designed to contain specific nucleotide substitutions as follows: 1) a shielding primer (GU4783) containing three silent mutations mutating the sgRNA and PAM sites targeted by the GU4788+GU4789 sgRNA (to prevent Cas9 binding the donor templates and the edited loci in the mutant parasites) as well as introducing a *BseYI* restriction site for restriction site fragment polymorphism (RFLP) analysis 2) Primer sets carrying the mutations of interest; V2721F mutation (GU4785) and V2752F (GU4784). A site directed mutagenesis of the cloned UBP-1 PCR product in the TOPO 2.1 vector was carried out using a QuikChange® multi-site directed mutagenesis kit (Agilent technologies) using the following primer combinations: GU4783+GU4784 for the V2752F single mutant and GU4783+GU4784+GU4785 for the double mutant. The resulting mutant fragments in the TOPO 2.1 vector were digested out and cloned into the linker site of the vector pG944 using the *HincII* restriction site to generate pG945 (single mutant) and pG946 (double mutant). For targeted mutation swapping and a second attempt to generate a double mutant line, a second sgRNA (GU5206+GU5207) upstream of the V2721F mutation was designed and cloned into the ABR099 vector as described. Donor DNA was amplified from the G1808^V2721F^ or pG946 vector to generate single or double mutation templates respectively using overlapping PCR as previously described (Heckman and Pease, 2007). Briefly, internal complementary primers (GU5190 and GU5191, supplementary Table 1) carrying 3 silent mutations (2 for mutating the sgRNA and PAM of the GU5206+GU5207 sgRNA, 1 to introduce the *SnaBI* restriction site for RFLP analysis) were used to amplify 2 overlapping PCR products from the G1808^V2721F^ DNA or pG946 plasmid upon linkage to *HincII* introducing outer primers GU5189 and GU4787 (supplementary Table 1). After gel purification, ~50ng of the overlapping PCR fragments were used as templates in a second round of PCR using the two outer primers (GU5189 and GU4787) to generate donor fragments with mutations of interest. The resulting fragments were subsequently cloned into the pG960 vector at the linker site using the *HincII* restriction site to generate the vectors pG963 (silent mutations to GU5206+GU5207 sgRNA, V2721F mutation) and pG962 (silent mutations to GU5206+GU5207 sgRNA, V2721F and V2752F mutation). All PCR reactions were carried out using the high fidelity KAPA hifi PCR kit (Roche). Plasmids were verified by Sanger DNA sequencing prior to further use.

### *P. berghei* animal infections

*P. berghei* parasites were maintained in female Theiler’s Original (TO) mice (Envigo) weighing between 25-30g. Parasite infections were established either by intraperitoneal injection (IP) of ~200µl of cryopreserved parasite stocks or intravenous injections (IVs) of purified schizonts. Monitoring of parasitaemia in infected mice was done by examining methanol fixed thin blood smears stained in Giemsa (Sigma) or flow cytometry analysis of infected blood stained with Hoescht 33342 (Invitrogen). Blood from infected mice was collected by cardiac puncture under terminal anaesthesia. All animal work was performed in compliance with UK home office licensing (Project reference: P6CA91811) and ethical approval from the University of Glasgow Animal Welfare and Ethical Review Body.

### Parasite lines and transfections

An 820 line that express green fluorescent protein (GFP) and red fluorescent protein (RFP) in male and female gametocytes respectively (Ponzi et al., 2009) was used for initial transfection experiments while the 1804cl1 line that constitutively express mCherry throughout the life cycle (Burda et al., 2015) was used for growth competition assays as a control. ~10µg of episomal plasmid DNA from the vectors described above was transfected by mixing with Nycodenz purified schizonts and electroporated using the Amaxa Nucleofector Device II program U-o33 as previously described (Philip et al., 2013). Parasites were then immediately IV injected into a tail vein of mice. Positive selection of transfected parasites was commenced 24 hours later by inclusion of pyrimethamine (Sigma) in drinking water.

### Genotype analysis of mutant lines

Blood was collected from parasite infected mice by cardiac puncture under terminal anaesthesia and lysed by resuspension in 1X E-lysis buffer (Thermo). Parasite genomic DNA was extracted using the Qiagen DNeasy Blood and Tissue kit according to manufactures’ instructions. Genotype analysis of the transfected or cloned parasite lines was analysed, initially by a dual PCR-RFLP. PCR using exterior primers (GU4894+GU4895 or GU5186+GU4895) was used to amplify fragments from the DNA of the mutant lines followed by restriction digests with either *BseYI* or *SnaBI* restriction enzymes to verify successful editing of the UBP-1 locus. Further confirmation of the mutations was carried out by Sanger DNA sequencing.

### *P. berghei in vitro* culture and drug susceptibility assays

For *in vitro* maintenance of *P. berghei*, cultures were maintained for one developmental cycle using a standardised schizont culture media containing RPMI1640 with 25mM hypoxanthine, 10mM sodium bicarbonate, 20 % foetal calf serum, 100U/ml Penicillin and 100μg/ml streptomycin. Culture flasks were gassed for 30 seconds with a special gas mix of 5% CO2, 5% O2, 90% N2 and incubated for 22-24 hours at 37°C with gentle shaking, conditions that allow for development of ring stage parasites to mature schizonts. Drug assays to determine *in vitro* growth inhibition during the intraerythroctic stage were performed in these standard short-term cultures as previously described (Franke-Fayard et al., 2008). Briefly, 1 ml of infected blood with a non-synchronous parasitaemia of 3-5% was collected from an infected mouse and cultured for 22-24 hours in 120 ml of schizont culture media. Schizonts were enriched from the cultures by Nycodenz density flotation as previously described (Philip et al., 2013) followed by immediate injection into a tail vein of a naive mouse. Upon IV injection of schizonts, they immediately rupture with resulting merozoites invading new red blood cells within minutes to obtain synchronous *in vivo* infection containing >90% rings and a parasitaemia of 1-2%. Blood was collected from the infected mice 2 hours post-injection and mixed with serially diluted drugs in schizont culture media in 96 well plates at a final haematocrit of 0.5% in a 200µl well volume. Plates were gassed and incubated overnight at 37°C. After 22-24 hours of incubation, schizont maturation was analysed by flow cytometry after staining the infected cells with DNA dye Hoechst-33258. Schizonts were gated and quantified based on fluorescence intensity on a BD FACSCelesta or a BD LSR Fortessa (BD Biosciences, USA). To determine growth inhibitions and calculate half-inhibitory concentrations (IC_50_), quantified schizonts in no drug controls were set to correspond to 100% with subsequent growth percentages in presence of drugs calculated accordingly. Dose response curves were plotted in Graph-pad Prism 7.

### *In vivo* drug assays

A modified Peters’ 4 day suppressive test was employed to assess i*n vivo* drug responses and or resistance profiles in the wildtype and mutant lines as previously described (Vega-Rodríguez et al., 2015). Parasitaemia was initiated by IP inoculation of between 10^6^-10^7^ parasites followed by three daily consecutive drug doses initiated ~4 hours post inoculation. CQ was prepared at 50mg/ml in 1X PBS and diluted to working stock in 1X PBS while ART was prepared at 12.5mg/ml in a 1:1 mixture of DMSO and Tween^®^ 80 (Sigma) followed by a ten-fold dilution in sterile water to an injectable working solution. All drugs were delivered by IP and were prepared fresh immediately before injection. Parasitaemia was monitored daily by flow cytometry and analysis of methanol fixed Giemsa stained smears.

### *In vivo* growth competition assays

Clonal mutant lines in the 820 background were mixed with the 1804cl1 line that constitutively express mCherry under the control of the *hsp70* promoter at a 1:1 mixture and injected intravenously in mice. Parasitaemia in the competition mixtures was quantified by flow cytometry quantification of mCherry positive parasites for the 1804cl1 proportional percentage and by subtracting the total parasitaemia (Hoescht positive) from the mCherry positive proportion for the 820 control and or mutant lines. Differentiation of the mCherry positive population from the RFP in the 820 line was carried out by applying flow compensation gating strategies (Supplementary Figure 3).

## Results

### CRISPR-Cas9 engineered mutations in UBP-1 select for ART and CQ resistance in *Plasmodium berghei in vivo*

To experimentally demonstrate that UBP-1 mutations confer ART resistance, we introduced *P. chabaudi* UBP-1 candidate mutation (V2697F and V2728F) equivalents (supplementary Figure 1) in the *P. berghei* 820 line using the CRISPR-Cas9 system developed and optimised in our lab (Figure 1A). Two plasmids were initially designed to either introduce the single mutation, V2752F (V2728F *P. chabaudi* equivalent) or both mutations, V2721F (V2697F *P. chabaudi* equivalent) and V2752F in an attempt to generate a double mutant (Figure 1A). Silent mutations to mutate the Cas9 cleavage site and introduce a restriction site *(BseYI*) were also introduced to prevent re-targeting of mutated loci by Cas9 for the former and diagnosis by RFLP for the latter (Figure 1A, 1B). Transfections of these plasmids into the 820 line yielded <1% mutants for the V2752F mutant line (G1807, pG945) and ~20% mutants for the V2721F and V2752F double mutant line (G1808, pG946) as confirmed by RFLP analysis (*BseYI* digestion) of the edited UBP-1 loci (Figure 1B). Since the efficiency was too low to clone out the mutant lines by serial dilution, we attempted a pre-emptive drug selection with CQ and ART of the G1807 and G1808 lines to examine if selective enrichment of the mutant population could be achieved. Indeed, after infecting mice with the G1808 line and treating for three consecutive days with ART at 20mg/kg, the recrudescent parasite population on Day 9 was enriched to >90% mutant population as confirmed by RFLP analysis (Figure 1C). Furthermore, CQ at 15mg/kg moderately enriched the same G1808 line, but not to the extent of ART challenge (Figure 1C). On the contrary, a very low level mutant enrichment of the G1807 line was observed with CQ at 15mg/kg while ART did not produce any enrichment in the same line. Interestingly, cloning of the G1808 ART enriched lines yielded six clones which were all single mutants positive for the V2721F mutation despite coming from a plasmid with donor templates that carried both the V2721F and V2752F mutations (Figure 1E). This suggests that the V2721F mutation specifically confers ART resistance, which explains the selective enrichment of this population upon ART pressure. These data also suggested that the double mutant parasites could either be lethal or unfit and henceforth outcompeted by single mutation carrying parasites in the presence of drug pressure. Sequence analysis of the G1808 line isolated after CQ challenge at 15mg/kg also revealed an enrichment for the V2721F mutation (supplementary Figure 2) suggesting that despite being principally enriched by ART, the V2721F mutation also modulates some low-level resistance to CQ. Meanwhile, when we challenged the G1807 line (V2752F single mutation) with CQ at a higher dose (30mg/kg), a recrudescent population on Day 10 was enriched to >60% (Figure 1D). Sanger sequencing of G1808 ART enriched and G1807 CQ enriched clones confirmed the presence of the single V2721F and V2752F mutations respectively, as well as the Cas9 cleavage silencing mutations and the silent mutations introducing the *BseYI* diagnostic restriction site (Figure 1F).

### The V2721F mutation confers observable phenotypic resistance to ARTs *in vivo* while the V2752F mutation confers high level resistance to CQ and low-level resistance to ARTs

We next quantitated the drug response profiles of the G1808^V2721F^ and G1807^V2752F^ cloned lines *in vitro* and *in vivo* using DHA, ART and CQ. In short term *P. berghei in vitro* drug assays, both the G1808^V2721F^ and G1807^V2752F^ parasites show no difference in sensitivity to DHA compared to the parental 820 line (Figure 2A, 2B). The lack of decreased drug sensitivity of both lines is consistent with the failure of the standard 72 hour drug assays to differentiate similar Kelch 13 ART resistant parasites from sensitive lines in *P. falciparum* (Dondorp et al., 2009, Witkowski et al., 2013). Meanwhile, a 1.8 fold increase in IC_50_ was observed for the G1807^V2752F^ line when challenged with CQ (Figure 2C) and not the G1808 ^V2721F^ (Figure 2D). However, rodent malaria parasites offer the advantage of experimental drug resistance assessment *in vivo*. This approach applied to G1808^V2721F^ demonstrated that this mutation does indeed confer resistance *in vivo* to ARTs compared to the parental 820 line. G1808^V2721F^ parasites survive three consecutive doses of 75mg/kg ART with the recrudescent population appearing on day 9 after last dosing while 820 wildtype parasites are effectively suppressed up to day 17 of follow-up (Figure 2E). Both the G1808^V2721F^ and 820 lines survive 45mg/kg dose of ART with the former having a slightly faster recrudescence rate on day 7 while the latter recrudesces a day later (Figure 2E). Both lines remain sensitive to 125mg/kg ART dose with no recrudescence observed up to day 17 (Figure 2E). In contrast, the G1807^V2752F^ line is highly resistant to CQ *in vivo* (Figure 2F), surviving three consecutive doses at 25mg/kg, with recrudescent parasites coming up on day 4 after the last dose as compared to the parental 820 line and the G1808^V2721F^ lines which are sensitive and are effectively suppressed up to day 17. Interestingly, the G1807^V2752F^ line also displays low level ART resistance at 75mg/kg dose, with parasites coming up on day 12, later than the G1808^V2721F^ line (Figure 2F). These data confirm that the V2721F mutation confers ART resistance while the V2752F mutation mediates resistance primarily to CQ and to some extent to ARTs. The recrudescence of the wildtype 820 and G1808^V2721F^ at 45mg/kg ART is also in agreement with our previous findings, that *P. berghei* is less sensitive to ARTs, especially in the spleen and bone marrow which could be the source of recrudescent infection at relatively lower doses (Lee et al., 2018).

**Figure 2:**
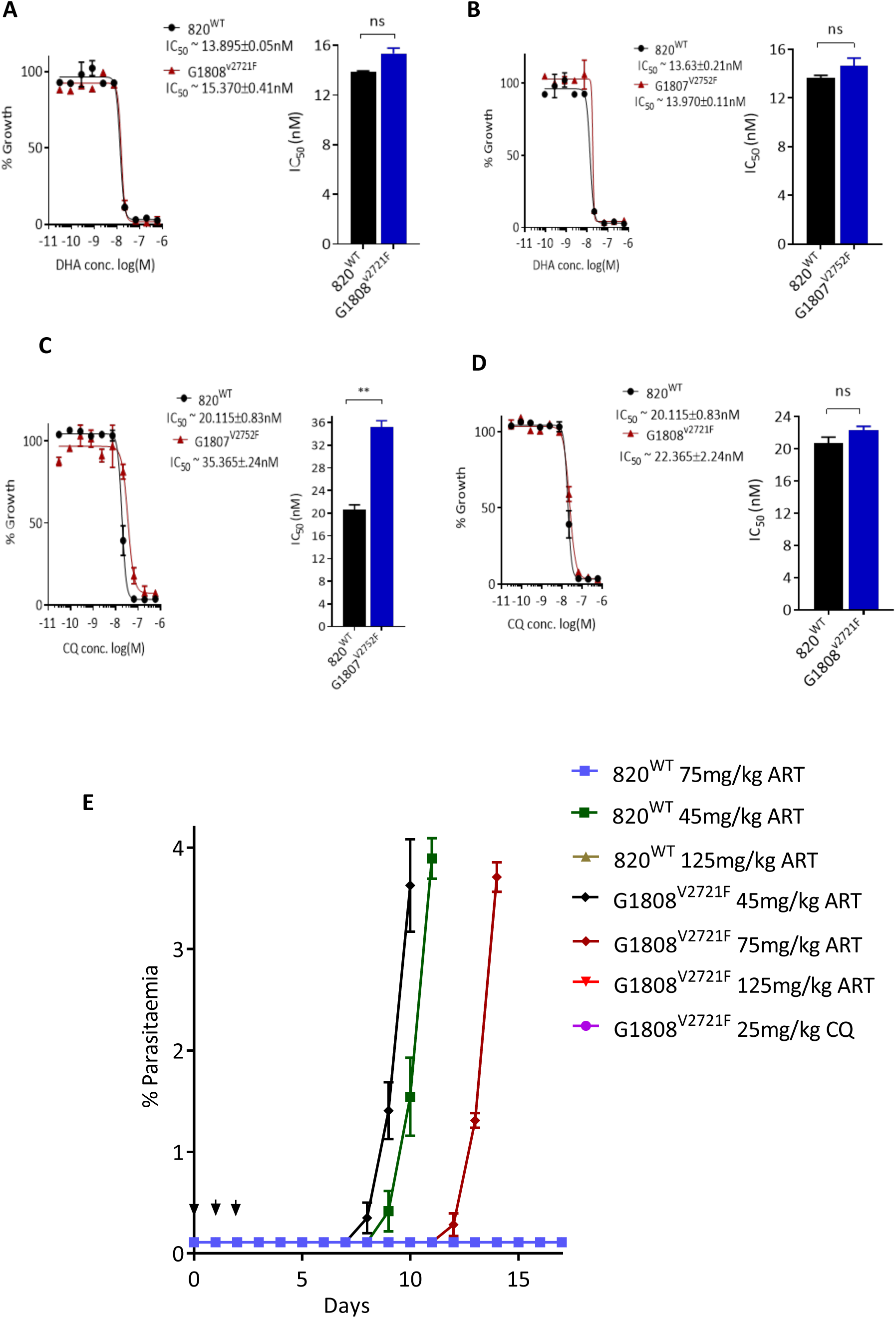
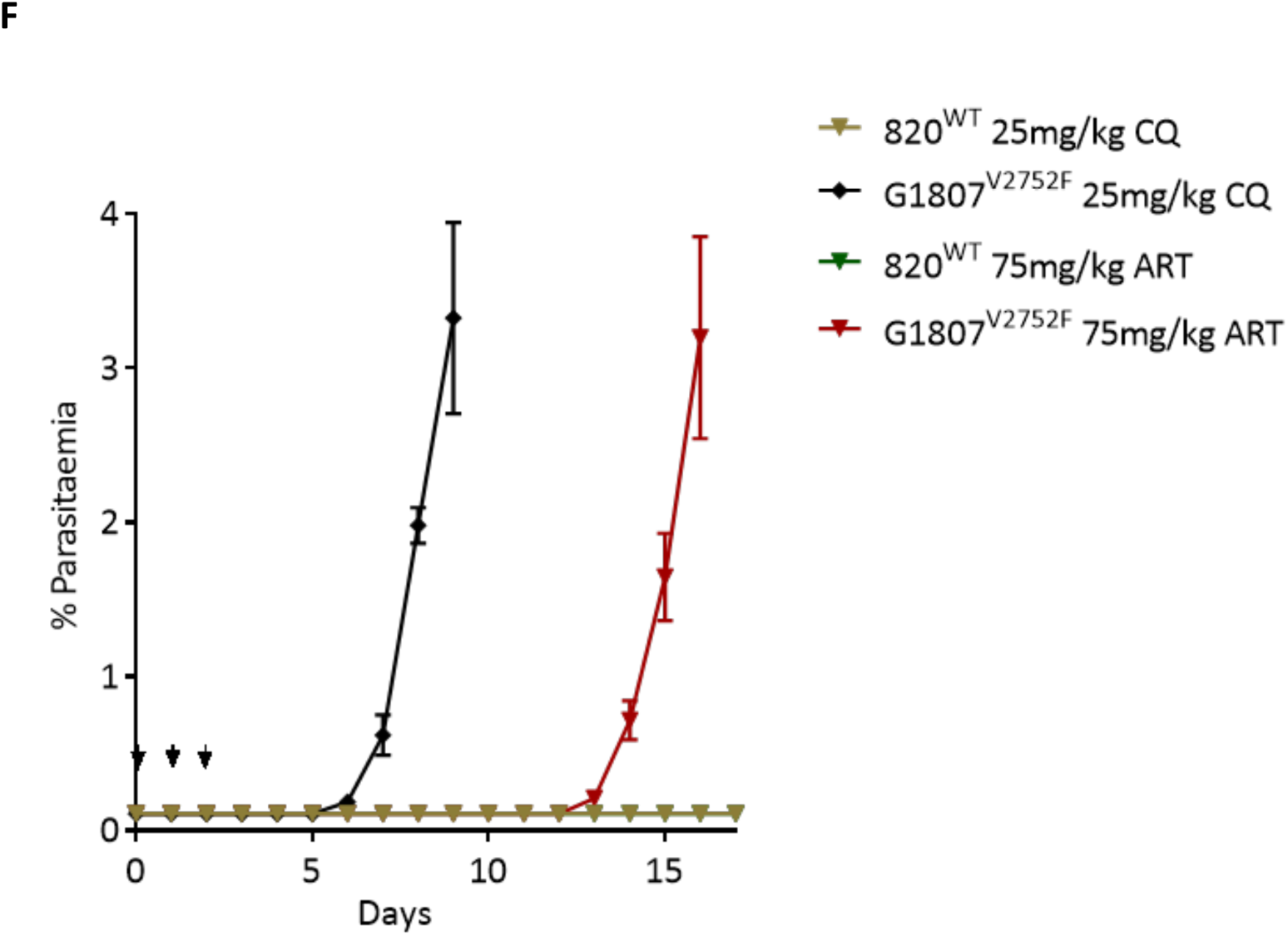
ART and CQ *in vitro* and *in vivo* resistance profiles of the G1807^V2752F^ and G1808^V2721F^ lines. DHA dose response curves and IC_50_ comparisons of the G1808^V2721F^ (**A**) and G1807^V2752F^ (**B**) lines relative to the wildtype 820 line. CQ dose response curves and IC_50_ comparisons of the G1807^V2752F^ (**C**) and G1808^V2721F^ (**D**) lines relative to the wildtype 820 line. Significant differences between mean IC_50_s or IC_50_ shifts were calculated using the paired t-test. Significance is indicated with asterisks; *p < 0.05, **p < 0.01, ***p < 0.001, ****p < 0.0001, ns; not significant. Modified Peters’ 4 day suppressive test to monitor resistance to ART and CQ *in vivo* in the G1808^V2721F^ (**E**) and the G1807^V2752F^ (**F**) mutant lines. Groups of three mice were infected with 1 × 10^6^ parasites on day 0. Treatment started ~1.5 hours later with indicated drug doses every 24 hours for three consecutive days (treatment days shown by arrows). Parasitaemia was monitored by microscopy analysis of Giemsa stained blood smears up to day 18. Error bars are mean parasitaemia from 3 mice groups.

### Growth of parasites carrying UPB-1 V2752F and V2721F mutations is impaired

The spread of drug resistance as is the case in most microbial pathogens is partly limited by detrimental fitness costs that accompany acquisition of such mutations in respective drug transporters, enzymes or essential cellular components. The G1807 and G1808 lines carrying UBP-1 V2721F and V2752F mutations respectively were each grown in competition with a parental line expressing mCherry *in vivo* and shown to be characteristically slow growing (Figure 3A-C). Comparatively, the G1807^V2752F^ mutation is severely impaired relative to the G1808^V2721F^ being completely outcompeted by day 8. These data and the earlier failure to generate the double mutant demonstrate that UBP-1 is an important (possibly essential) protein for parasite growth and that acquisition of resistance through mutation of UBP-1 confers mutation specific fitness costs.

**Figure 3:**
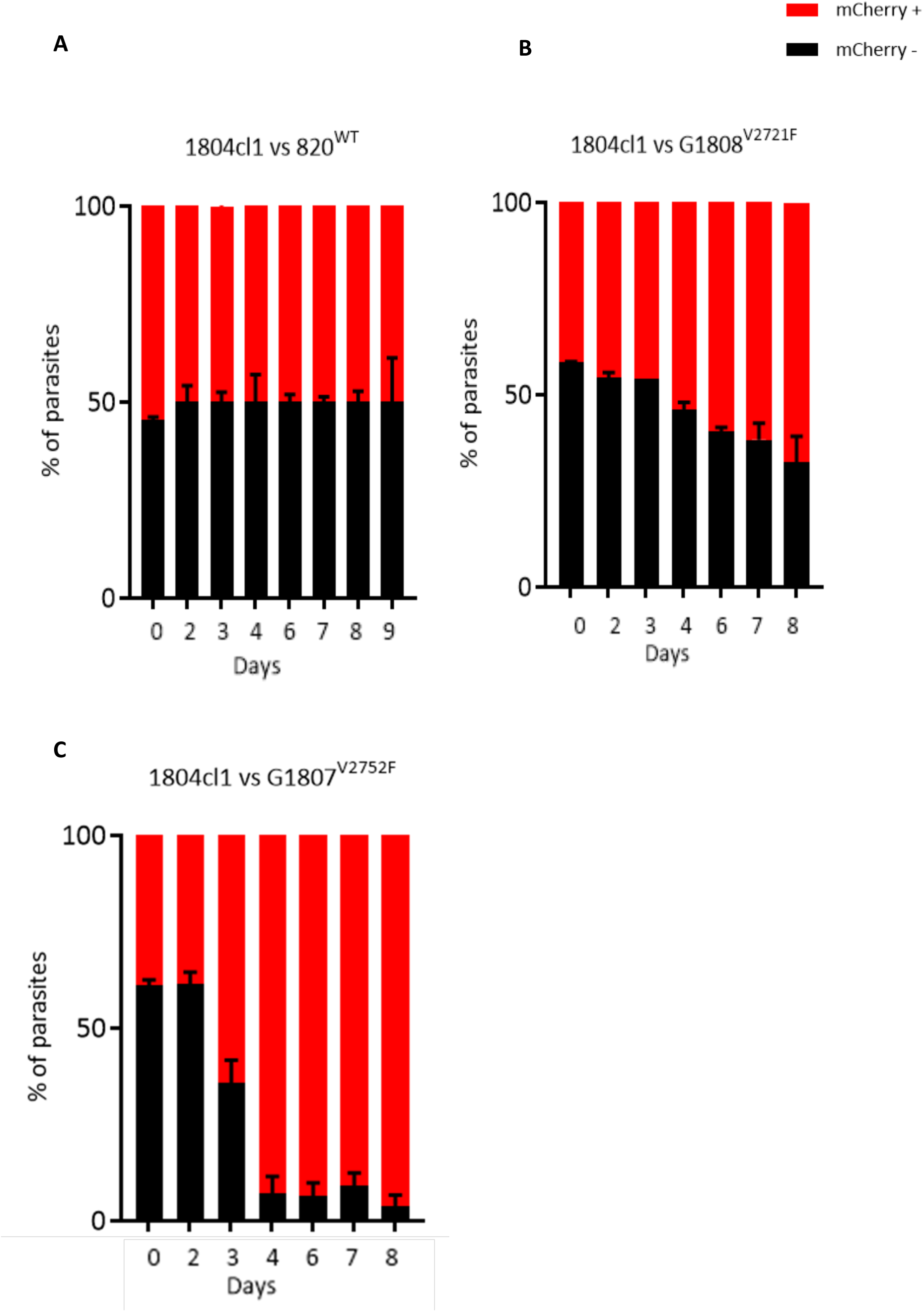
Growth kinetics of the 820, G1808^V2721F^ and G1808^V2752F^ relative to the 1804cl1 line. The 1804cl1 line constitutively express mCherry under the control of the *hsp70* promoter. The 820, G1808^V2721F^ and G1808^V2752F^ were mixed with the 1804cl1 at a 1:1 ratio and injected at a parasitaemia of 0.01% intravenously on Day 0. Daily percentages of representative parasitaemia of the 820 or mutant lines in the competition mixture were quantified by subtracting the total parasitaemia based on positivity for Hoescht DNA stain from the fraction of the population that is mCherry positive (1804cl1) as determined by flow cytometry. On day 4, when parasitaemia was ~5%, blood from each mouse was passaged into new naïve host and parasitaemia was monitored until day 8. Percentage population changes of the mutant and wildtype lines relative to the 1804cl1 in the 820 (**A**), G1808^V2721F^ (**B**) and G1807^V2752F^ (**C**). Error bars are standard deviations from three biological repeats.

### Reversal of the V2752F mutation restores CQ sensitivity in the G1807^V2752F^ line while introduction of the V2721F in the same line appears to be lethal

Drug pressure can select in the long or short term for mutations in sensitive parasite populations that would affect responses to the same drug. To further confirm that the resistance phenotypes observed in our mutant lines were due to the V2721F or V2752F mutations and not possible secondary mutations which may have been acquired during the pre-emptive drug pressure, we attempted to reverse the V2752F mutation to see if wildtype drug phenotypes can be restored in the G1807^V2752F^ line. Using a CRISPR Cas9 editing strategy similar to the one outlined above, a sgRNA targeting a region ~50bp upstream of the V2721F mutation was designed and cloned in the Cas9 expressing vectors (Figure 4A). ~710bp of donor DNA (GU5189 + GU4787) containing the V2721F (for targeted mutation swap) or both the V2721F and V2752F mutations (for a forced introduction of the V2721F in the G1807^V2752F^ background) was used to generate the vectors pG963 and pG962 respectively (Figure 4A). Silent mutations mutating the PAM site as well as introducing a second restriction site, *SnaBI*, for RFLP analysis were also included. Transfection of the G1807^V2752F^ line with pG963 and pG963 vectors successfully edited the UBP-1 loci generating the G1918 and G1919 lines respectively with >85% efficiency as confirmed by *SnaBI* RFLP analysis (Figure 4A). Cloning and sequencing of the G1918 lines revealed successful targeted mutation swap, introducing the V2721F mutation and re-editing of the 2752F to 2752V wildtype genotype (Figure 4B, 4C). Phenotype analysis of the G1918 clone line revealed a similar ART resistance profile to ART at 75mg/kg as the G1808^V2721F^ line while CQ sensitivity was completely restored (Figure 4D). This provided further experimental evidence, that the resistance profiles observed were due to the V2721F or V2752F amino acid substitutions and not the introduced silent mutations or secondary mutations that may have been acquired during the pre-emptive drug exposure. Interestingly, cloning and sequencing of the G1919 (Figure 4B, 4E) line revealed successful introduction of the silent mutations (PAM mutating and *SnaBI*) while the V2721F mutation was absent in all four clonal lines yet retained the parental V2752F mutation. This suggested that introduction of the V2721F in the V2752F background is lethal to the parasite and supported our failed attempts to generate the double mutant line (Figure 1). Indeed, detailed sequence analysis of the transfected parasite populations before cloning revealed the presence of only one mutation trace in either the G1808 or G1919 lines (despite the donor DNA containing both mutations) confirming that the double mutant parasites do not survive or are severely growth-impaired and quickly overgrown by the single mutation parasites (Figure 4F, 4G).

**Figure 4:**
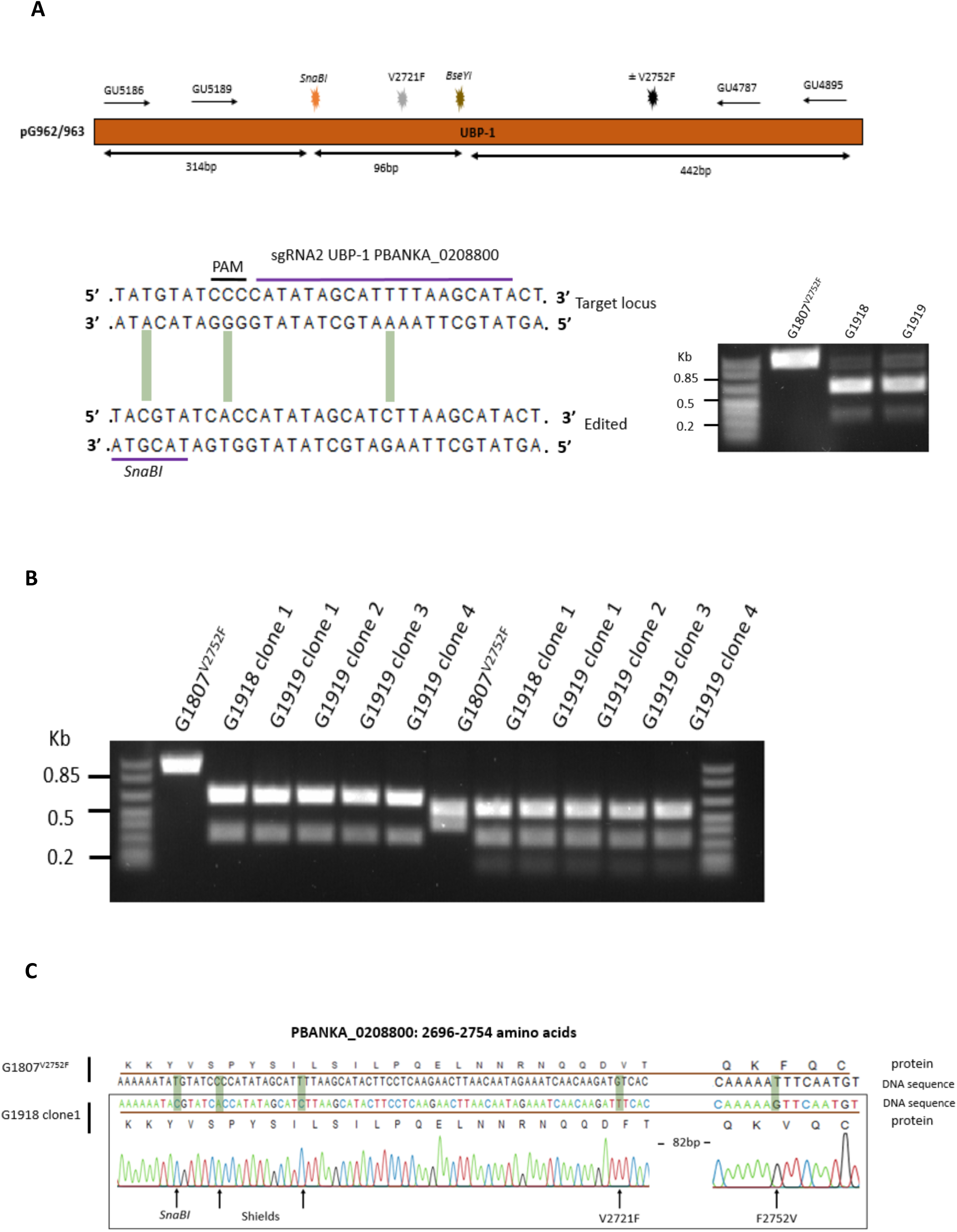
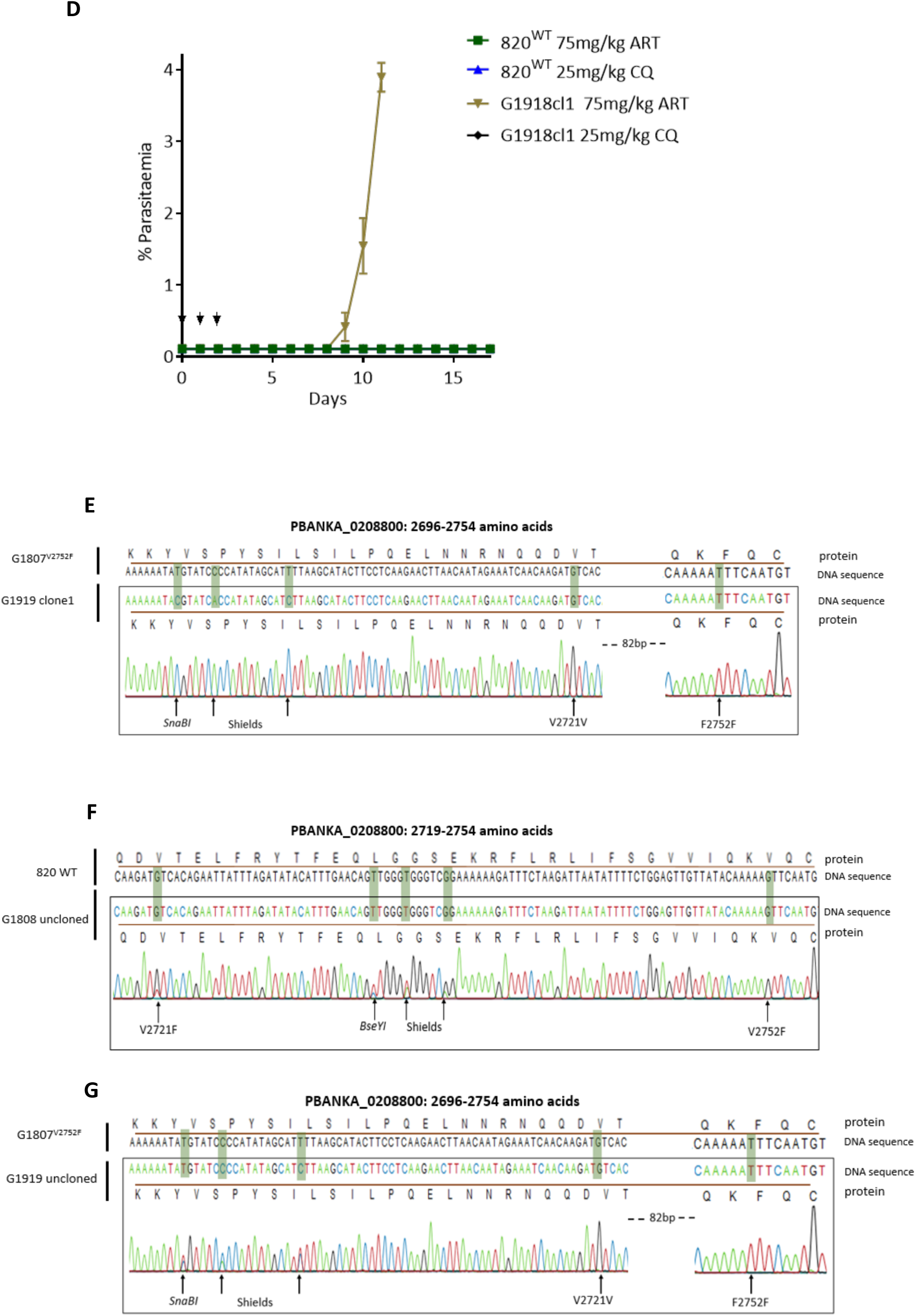
Swapping of the V2752F to V2721F mutations and attempted generation of a double mutant in the G1807^V2752F^ line. **A.** Schematic of the UBP-1 donor DNA in the pG962 and pG963 vectors, a 20bp guide RNA used to target the UBP-1 region upstream of the V2721F mutation in the Cas9 expressing vectors with introduced silent mutation sites indicated, RFLP (*SnaBI* digestion) analysis of PCR products (GU5186 + GU4895) of the G1918 and G1919 lines. **B.** RFLP analysis of the cloned G1918 and G1919 lines. First six lanes to the left are RFLP analyses of G1918 and G1919 cloned lines PCR products digested by *SnaBI* including the parental G1807^V2752F^ line, while 6 lanes to the right are the same clones digested by both *SnaBI* and *BseYI*. **C.** Sequencing of G1918 clone1 showing successful swapping of the V2752F in the parent G1807^V2752F^ line to the V2721F mutation. **D.** Phenotype analysis of the G1918 clone1 line showing resistance to ARTs at 75mg/kg and complete restoration of CQ sensitivity. Sequence analysis of the G1919 clone 1 (**E**) line and the G1808 (**F**) and G1919 (**G**) uncloned lines showing absence of double mutant populations.

## Discussion

Ubiquitin hydrolases or deubiquitinating enzymes (DUBs) are essential elements of the eukaryotic ubiquitin proteasome system (UPS) which is primarily involved in maintaining cellular protein homeostasis and responding to stress. Despite the proposed involvement of *Plasmodium* DUBs in modulating resistance to multiple drugs, lack of conclusive experimental evidence has thus far limited studies into their detailed involvement in mode of action and or resistance phenotypes such as those observed with ARTs. In this study, using a CRISPR-Cas9 mediated reverse genetics approach; we have provided experimental evidence on the direct involvement of a DUB (UBP-1) in mediating resistance to ART and CQ, more importantly under *in vivo* conditions. As the debate into the mechanism of action and resistance to ARTs continues, a consensus understanding is converging that ART resistance is more complex as several factors, genetic determinants and possibly mechanisms of action appear to be involved. In *P. falciparum*, ART resistance is confined to early ring stage parasites which has been translated in laboratory conditions to increased survival in ring stage survival assays (Witkowski et al., 2013). Mutations in Pfkelch13, PfCoronin as well as transient (hypo-hyperthermic) temperatures have all been shown to enhance ring stage parasite survival in the RSAs (Henrici et al., 2019b, Straimer et al., 2015, Demas et al., 2018). As demonstrated in this study, ART and CQ resistance can also be mediated by mutations in UBP-1 underscoring a mechanism of cross-resistance and some commonality in mode of action between CQ and ART especially relating to haemoglobin digestion and trafficking in malaria parasites (Klonis et al., 2011).

The UBP-1 V2728F mutation was previously designated as a principle determinant of ART resistance despite its common fixation with mefloquine and higher doses of CQ (Hunt et al., 2010). Contrary to this argument, ART did not enrich for this mutation (V2752F) in our study enriching for the V2721F mutation instead which was fixed with artesunate in *P. chabaudi*. However, enrichment of the V2752F mutation with a higher dose of CQ was achieved showing that this mutation does indeed modulate resistance to CQ while the V2721F mutation is chiefly responsible for the ART resistance phenotype in the *P. berghei* model *in vivo*. Interestingly, drug challenge of these mutant lines *in vivo* revealed that both mutations also give low-level cross-resistance to ARTs and CQ. This confirms that both of these UBP-1 mutations modulate some form of resistance to both ARTs and CQ albeit to some differing degrees which is, therefore, in strong agreement with previous observations in *P. chabaudi* (Hunt et al., 2010). This also demonstrates a plurality of pathways to resistance involving the same target. Recently, the exact equivalent UBP-1 mutations in *P. falciparum*, V3275F and V3306F have been successfully engineered (Henrici et al., 2019a). In *P. falciparum* UBP-1, the V3275F mutation (V2721F *P. berghei* equivalent) shows enhanced survival to DHA in RSAs but remains sensitive to CQ. However, unlike in *P. berghei*, the V3306F (V2752F *P. berghei* equivalent) showed no enhanced survival to DHA in RSAs or resistance to CQ (Henrici et al., 2019a). Whilst not entirely in agreement with the data reported here, this could be due to limitations in the ability of *in vitro* assays to fully predict actual drug responses *in vivo* which these data highlight and has been the concern with Kelch 13 mutations recently (Sa et al., 2018). These observations may also somewhat be confounded by species specific differences in drug responses, pharmacodynamics, modes of action and resistance that in part remain to be fully investigated. For example, previous and original linkage studies in *P. chabaudi* identified additional mutations in an amino acid transporter (*pcaat*) as being strongly associated with CQ resistance phenotypes in tandem with UBP-1 mutations (Hunt et al., 2010). Even though this could partly explain the observed *in vitro* sensitivity of *P. falciparum* V3275F mutants to CQ, our data suggests that UBP-1 mutations are sufficient to mediate resistance to both ARTs and CQ as the reversal of the V2752F mutation performed in this study, for example, completely restores observable CQ sensitivity. This has provided, therefore, additional independent evidence on the direct causative role of UBP-1 mutations in mediating resistance not just to ARTs, but CQ as well. The study also illustrates the potential of the *P. berghei* rodent model in proving causality to antimalarial drug resistance phenotypes under *in vivo* conditions especially in light of recent reported discrepancies between some *in vitro* RSA resistance profiles of *P. falciparum* Kelch 13 mutants and actual *in vivo* phenotypes using the Autos monkey model (Sa et al., 2018).

Interestingly, the V2721F and V2752F mutation carrying parasites are characteristically slow growing and are easily outcompeted in the presence of non-mutants. Natural *P. falciparum* UBP-1 mutations have been reportedly associated with ART treatment failure in Kenya (Henriques et al., 2014, Borrmann et al., 2013), SEA (Cerqueira et al., 2017) and more recently in Ghana (Adams et al., 2018) (Supplementary Figure 4). However, unlike their rodent counterparts which associate with ART resistance, the natural reported E1528D and D1525E mutations occur towards the less conserved N-terminus of the protein and outwith the conserved, bioinformatically predicted UBP-1 catalytic domain (Hunt et al., 2007) (supplementary Figure 1). This would suggest that acquisition of the mutations at the well conserved C-terminal in *P. falciparum* has a potential growth defect as we have observed with *P. berghei* in this study. However, as these upstream mutations are not conserved between *P. falciparum* and *P. berghei* UBP-1, we cannot test the hypothesis in this model. In fact, *P. falciparum* UBP-1 is highly polymorphic with over 480 reported single nucleotide polymorphisms (SNPs) https://plasmodb.org all of which are in the N terminal region. PfUBP-1 has also been recently shown to be undergoing a strong positive selection in SEA (Ye et al., 2019). UBP-1 mutations could, therefore, be an independent avenue to which ART or multidrug resistance phenotypes could emerge in endemic regions like has been seen in Africa (Ghana and Kenya), without actually requiring a permissive genetic background as seems to be the current landscape with Kelch 13 mutations. However, there are constraints upon the evolution of drug resistance and UBP-1. Whilst these data confirm that a single protein that does not transport drugs can mediate resistance to two quite distinct drug entities, it was not possible to generate a *P. berghei* line that simultaneously contained the two UBP-1 drug resistance mutations examined in this study.

In yeast, UBP-1 localises to the endoplasmic reticulum playing roles in protein transport specifically internalisation of substrates across membranes (Schmitz et al., 2005). Mutations in UBP-1 could, therefore, modulate endocytosis of important essential host derived products such as haemoglobin to the digestive vacuole in a similar manner thereby reducing exposure of the parasite to activated drug for both ARTs and CQ. Interestingly, mutations in the AP2 adaptor complex that is involved in clathrin mediated endocytosis have also been implicated in ART resistance in rodent malaria parasites (Henriques et al., 2013). One of the AP2 adaptor complex mutation (I592T) has been recently engineered in *P. falciparum* and has been shown to enhance ring stage parasite survival in RSAs (Henrici et al., 2019a). This further suggests that inhibition of the endocytic trafficking system is a possible generic mechanism for the parasites to survive lethal doses of drugs that require transport and activation in the digestive vacuole. This would further explain the multidrug resistance phenotype observed with the UBP-1 mutations in *P. chabaudi* and *P. berghei* in this study.

Acquisition of the V2728F mutation in *P. chabaudi* was structurally predicted to reduce deubiquitination (Hunt et al., 2007). In such a situation, the cellular increase in ubiquitinated proteins would be anticipated to positively feedback to the cellular machinery to rapidly degrade protein substrates at the 20s proteasome promoting a non-specific and rapid protein turnover or impaired substrate trafficking. This would result in generally slow growing parasite with reduced expression of, for example, multi-drug resistance transporters as well as reduced endocytosis of host-derived products like haemoglobin, which would in turn modulate parasite responses to these drugs. More recently, functional studies have revealed that PfKelch13 (known determinants of ART resistance) localises to the parasite cytosomes and play a role in haemoglobin trafficking (Yang et al., 2019). Consequently, PfKelch13 mutations have been shown to lead to a partial loss of PfKelch13 protein function leading to decreased haemoglobin trafficking to the parasite digestive vacuole and less DHA activation which in turn mediates parasite survival (Yang et al., 2019). This is indeed in agreement with our hypothesis on the consequences of UBP-1 mutations which in a similar manner could impair trafficking of haemoglobin leading to less activation of ARTs and CQ. The experimental validation of these mutations in ART and CQ resistance, therefore, provide an additional understanding of drug resistance in malaria parasites. Further work, to identify and characterise proteins and pathways which interact with UBP-1 in modulating ART and CQ resistance will hopefully further illuminate the unique involvement of the parasite UPS network in ART and CQ resistance possibly through direct comparison with Kelch 13 mutation equivalents. Furthermore, the *P. berghei* model provides a useful system in which to investigate the interplay and impact of simultaneous mutations of both K13 and UBP-1 *in vivo* as well as assess whether PfKelch13 mutations would modulate resistance to CQ under *in vivo* conditions.

In conclusion, the work presented here provides direct experimental evidence for the involvement of conserved mutations in a polymorphic ubiquitin hydrolase protein that serves as a nexus for resistance to two very diverse classes of drugs. The findings also underscore the potential difficulties that *in vitro* assays may have in appropriately assigning mutant parasites with appropriate phenotypes in absence of conclusive *in vivo* measurements. *P. berghei* should therefore, be a suitable and adaptable *in vivo* model for the rapid evaluation and/or genetic engineering of mutations associated with human-infectious *Plasmodium* drug resistance observed in the field for concurrent assigning of drug resistance phenotypes under both *in vitro* and *in vivo* conditions.

## Acknowledgments

We would like to thank Mathias Matti for meaningful discussions; Diane Vaughan and the iii flow cytometry facility for assistance. This work was supported by grants from the Wellcome Trust to A.P.W (083811/Z/07/Z; 107046/Z/15/Z). M.P.B. is funded by a Wellcome Trust core grant to the Wellcome Centre for Integrative Parasitology (104111/Z/14/Z). N.V.S is a Commonwealth Doctoral Scholar (MWCS-2017-789), funded by the UK government.

## Author contributions

N.V.S conceived the experiments, performed data curation, analysis, investigation, validation, visualisation and writing of original draft. K.R.H, A.B.R, M.T.R and M.P.B participated in formal data analysis, investigation, validation, review and editing. A.P.W conceived the study, experiments, analysis, investigation, validation, writing of original draft, review, editing and supervision.

---

**Supplementary table 1:**
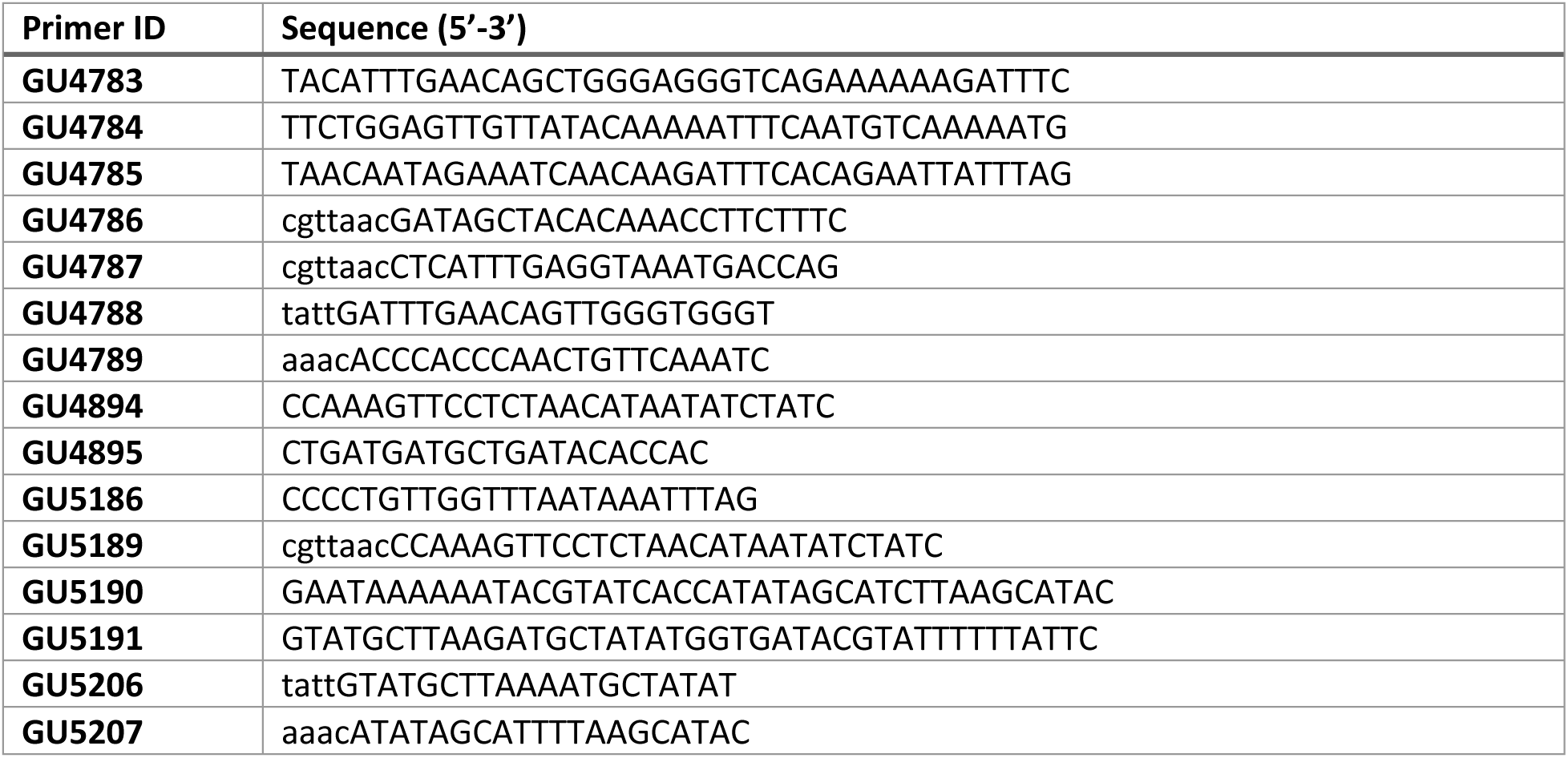
List of primers used

**Supplementary Figure 1:**
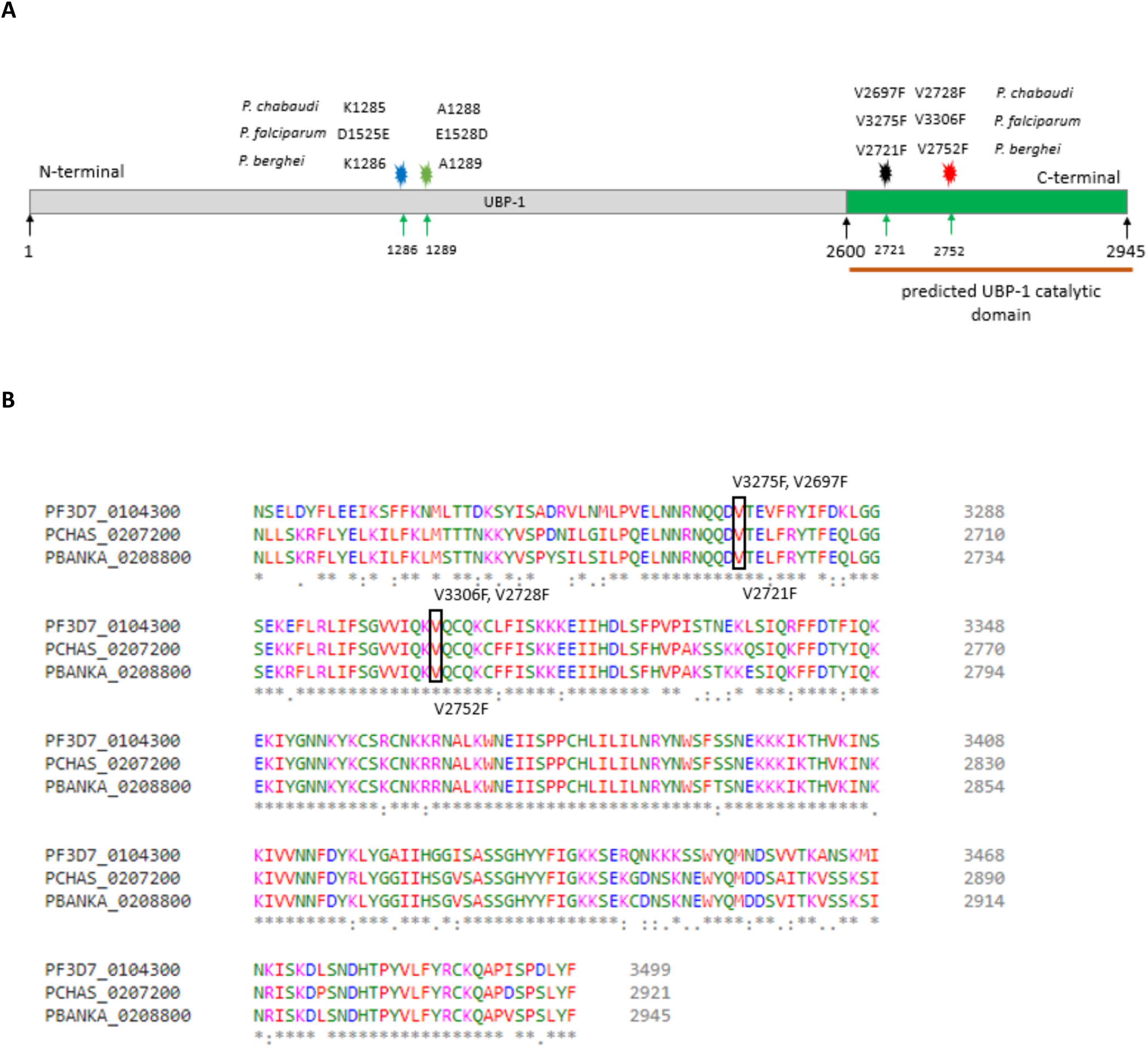
Cartoon of *P. berghei* UBP-1 and sequence alignment of *P. falciparum*, *P. chabaudi* and *P. berghei* at the C-terminal. **A.** *P. berghei* UBP-1 showing the predicted catalytic domain and localisation of the engineered mutations and their *P. falciparum P. chabaudi* equivalents. Positions of *P. falciparum* UBP-1 D1525E and E1528D mutations which have been reported in the field but are not conserved in *P. berghei* and *P. chabaudi* are indicated. **B**. Sequence alignment of *P. falciparum*, *P. chabaudi* and *P. berghei* at the conserved C-terminal. Mutation sites are indicated for *P. falciparum* and *P. chabaudi* on top and *P. berghei* on the bottom. Conserved sites are indicated by the * symbol.

**Supplementary Figure 2:**
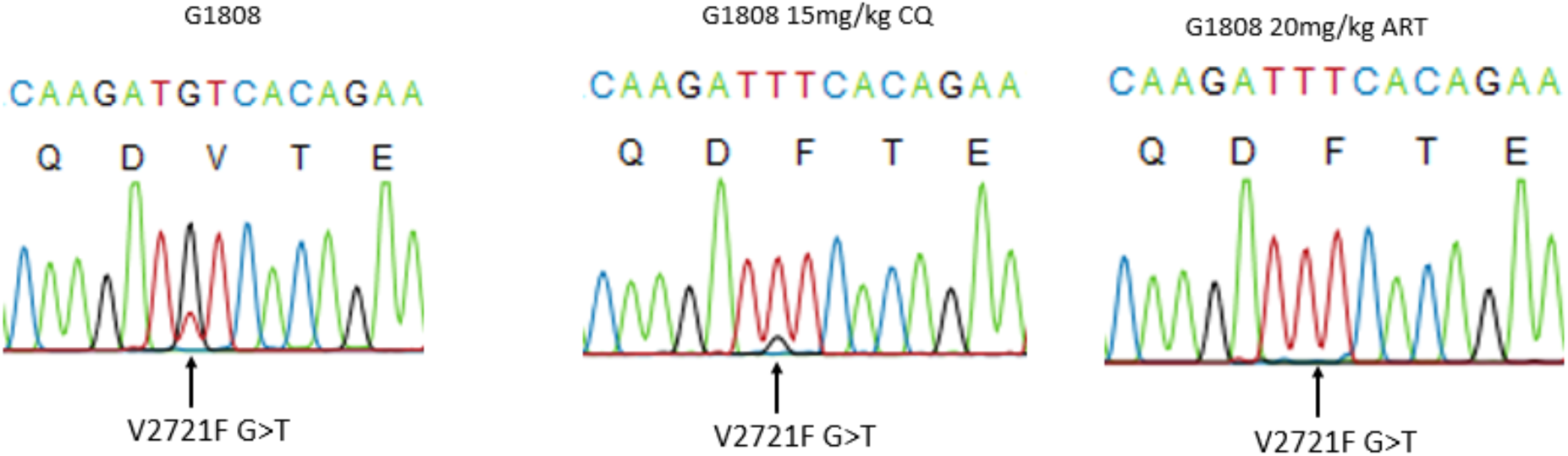
DNA sequencing and trace analysis of CQ and artemisinin challenged G1808 lines (Figure 1C). Enrichment of the V2721F mutation by CQ is observable despite the absence of *in vitro* and *in vivo* phenotypes in clonal lines carrying the same mutation.

**Supplementary Figure 3:**
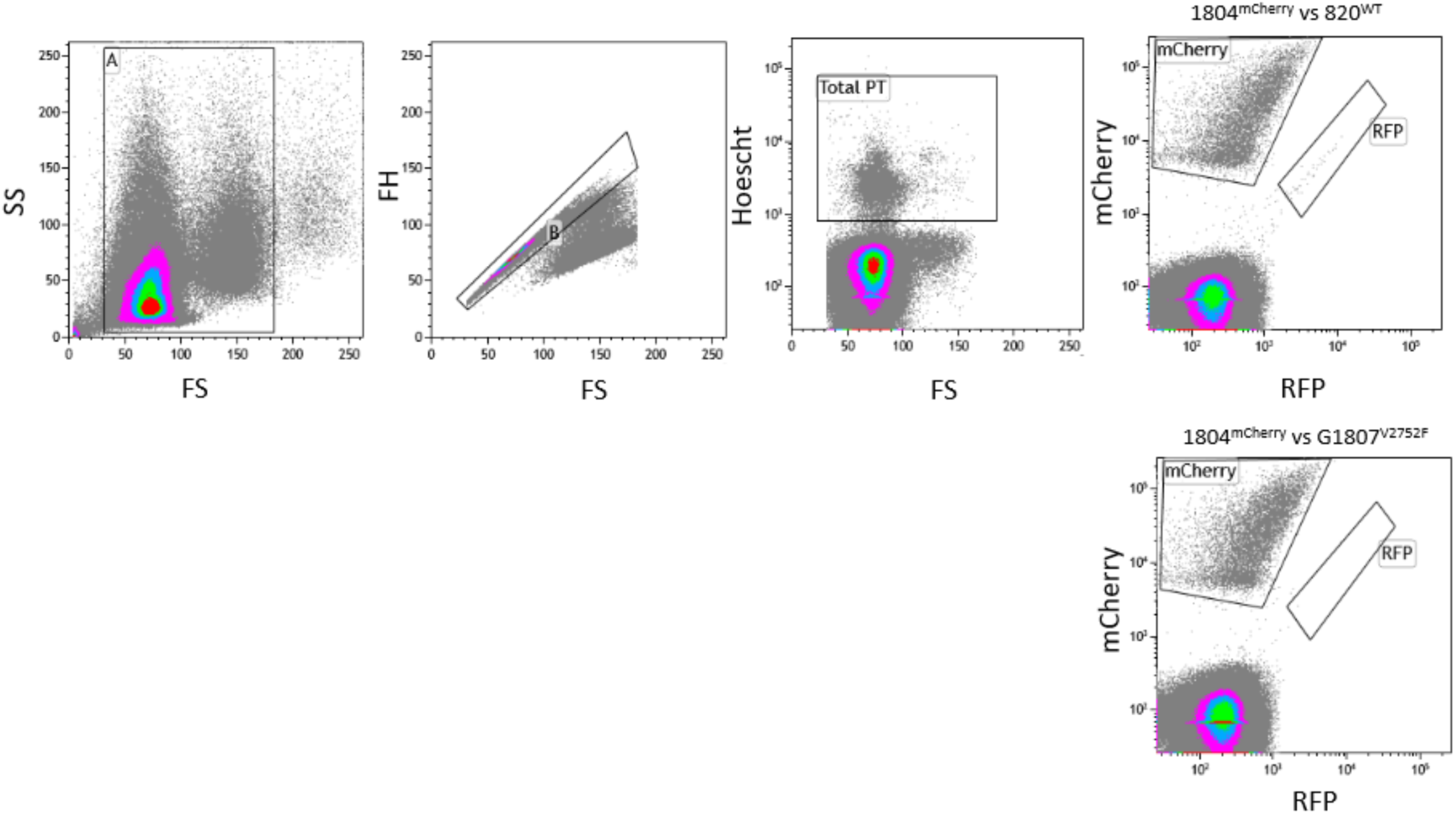
Flow cytometry gating strategy for growth competition experiments. Representative flow cytometry gating strategies for growth competitions of wildtype and mutant UBP-1 lines on Day 3. Acquired events were plotted on a forward (FS) and side (SS) scatter. Gate A was drawn to exclude debris. Events from gate A were plotted on a FS vs FH where gate B was drawn to exclude cell clumps and potential doublets. Events from gate B were then plotted on Hoescht vs FS and Total PT gate was drawn to quantify total parasitaemia. Events from gate B were also plotted on mCherry vs RFP where the mCherry positive population was distinguished from RFP positive female gametocytes by applying compensation spill-over filters that allow discrimination of the two colours as illustrated in the plots. Parasitaemia of mutant parasites was quantified by subtracting the mCherry positive population from the total parasitaemia as quantified by Hoescht staining of parasite DNA.

**Supplementary Figure 4:**
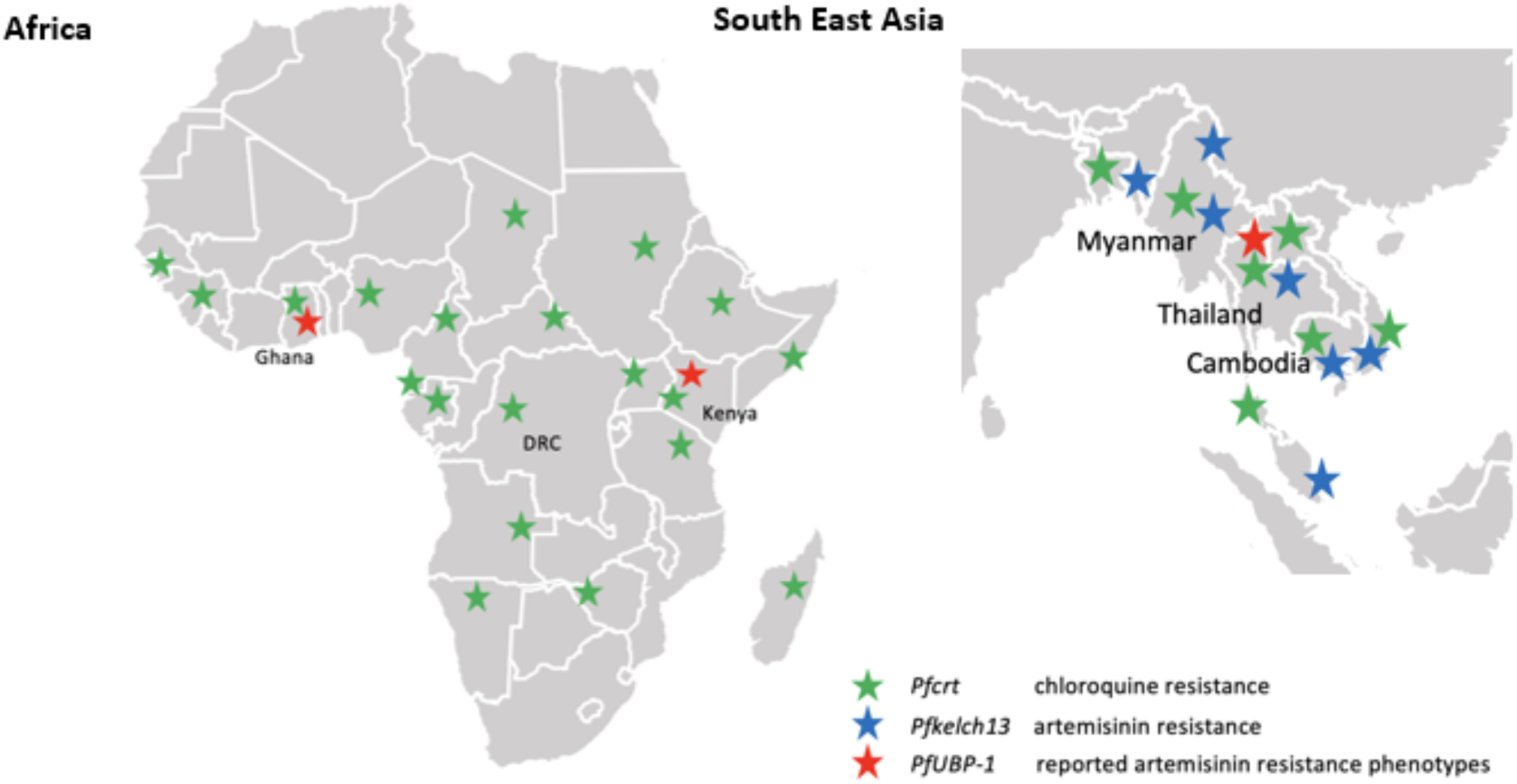
Distribution of ART and CQ resistance mutations in Africa and South East Asia. CQ resistance is believed to have originated in SEA and some parts of South America and eventually spread to Africa (Ecker et al., 2012). Current distribution of Kelch 13 mutations in SEA (Mbengue et al., 2015, Ashley et al., 2014, Menard et al., 2016) and reported UBP-1 polymorphisms (Cerqueira et al., 2017, Adams et al., 2018, Henriques et al., 2014, Borrmann et al., 2013).

